# SLAE: Strictly Local All-atom Environment for Protein Representation

**DOI:** 10.1101/2025.10.03.680398

**Authors:** Yilin Chen, Tianyu Lu, Cizhang Zhao, Hannah K. Wayment-Steele, Po-Ssu Huang

**Affiliations:** Stanford University, Department of Bioengineering; University of Wisconsin–Madison, Department of Biochemistry

## Abstract

Building physically grounded protein representations is central to computational biology, yet most existing approaches rely on sequence-pretrained language models or backbone-only graphs that overlook side-chain geometry and chemical detail. We present SLAE, a unified all-atom framework for learning protein representations from each residue’s local atomic neighborhood using only atom types and interatomic geometries. To encourage expressive feature extraction, we introduce a novel multi-task autoencoder objective that combines coordinate reconstruction, sequence recovery, and energy regression. SLAE reconstructs allatom structures with high fidelity from latent residue environments and achieves state-of-the-art performance across diverse downstream tasks via transfer learning. SLAE’s latent space is chemically informative and environmentally sensitive, enabling quantitative assessment of structural qualities and smooth interpolation between conformations at all-atom resolution.

## 1 Introduction

Proteins are the fundamental machinery of life, carrying out processes from catalysis and signaling to structural organization. Their remarkable functional diversity arises not only from their amino acid sequences but from the intricate three-dimensional structures into which those sequences fold.

Within protein structures, the backbone and side chain atoms act as an intricately coupled system that establishes local atomic environments through hydrophobic packing, hydrogen-bonding networks, and electrostatic interactions. These residue-level environments mediate conformational preferences and side chain dynamics, linking the global fold to the specific interactions that underlie protein function. Representing these interactions in a concise, learnable form is therefore essential for generalizable and physically grounded models of protein structure and function.

Current representations, either through protein language model(PLM) or sequence-structure joint embedding, lack the ability to isolate physical interactions from evolutionary information, and often needed to adopt backbone-only structure info to reduce computational demands. As a result, the field remains limited by the absence of a general-purpose pretraining framework that extracts, compresses, and transfers knowledge of all-atom structure across proteins and downstream applications.

We propose **SLAE** (**S**trictly **L**ocal **A**ll-atom **E**nvironment autoencoder), a framework for protein representation learning that models a protein as a set of residue-centric chemical environments. To promote generalizability and a physically grounded view, SLAE enforces an informational bottleneck by restricting the encoder to strictly local atom graphs and pair it with an asymmetric decoder that must recover full structure. When this reconstruction task is solved, the resulting tokenization of structure emerges jointly from the representation and the model, emphasizing physically meaningful interactions rather than heuristic features. Fully connected local atom graphs capture interactions between a residue and its neighboring atoms and are computationally tractable during pretraining. We show these local representations are sufficient to reconstruct all-atom Cartesian coordinates with high fidelity.

We design an all-atom autoencoder architecture that separates local and global reasoning across the encoding and decoding stages. An SE(3)-equivariant graph encoder maps each local environment to a rotation/translation-invariant residue token. A Transformer decoder with self-attention then aggregates these tokens to model long-range couplings and reconstruct coherent global geometry. This residue-level bottleneck forces the encoder to distill the packing signals such as covalent bonds, hydrogen-bond motifs, and steric/electrostatic cues that the global decoder requires to reconstruct long-range geometry, facilitating transfer across tasks. We introduce a physics-augmented pretraining objective that couples self-supervised (i) all-atom coordinate reconstruction, (ii) sequence recovery, and supervised (iii) Rosetta-derived inter-residue energies. These complementary signals act as a multi-view regularizer, aligning the latent space with atomistic structure, biochemical signal and energetics, yielding embeddings that vary smoothly with conformation and are interpretable along axes of side-chain chemistry, solvent exposure, and secondary structure.

SLAE supports multiscale readouts: atom and residue embeddings for fine-grained local characterization, and pooled protein-level features for global structure. This flexibility allows downstream task heads to focus on single residues, interfaces, or entire folds using a single pretrained representation. We demonstrate that pretraining directly on all-atom protein structures yields features that transfer effectively. Across benchmarks on multiple resolution scale tasks including fold classification, protein–protein binding affinity, single-point mutation stability, and NMR chemical shifts, SLAE achieves state-of-the-art or on-par performance.

### Main contributions

With the SLAE framework, we **(i)** propose a residue-centered, local atom-graph protein representation, and show it is sufficient for high-fidelity all-atom reconstruction; **(ii)** propose the energy regression task for reconstruction pretraining guidance; **(iii)** design local encoding and global decoding stages in all-atom autoencoder to encourage compact and transferable residue embeddings; **(iv)** achieve state-of-the-art on diverse downstream tasks with transfer learning; **(v)** show that the above design allow an interpretable latent space.

## 2 Related Work

### Protein Representation Pretraining

Protein representation learning has followed two main tracks. *Sequence pretraining* with protein language models (PLMs) on massive corpora captures evolutionary constraints but lacks explicit structure information (Meier et al., 2021; Lin et al., 2023). In parallel, graph denoising objectives noises sequence or structural features and train graph models to recover them (Zaidi et al., 2022; Jamasb et al., 2024), capturing global context while abstracting away side-chain geometry. Neither paradigm learns atomistic features as the primary signal. SLAE departs by *pretraining directly on all-atom coordinates reconstruction* and showing that features learned from atomistic geometry are sufficient for high-fidelity coordinate reconstruction and downstream transfer.

Sequence-structure co-embedding approaches pair PLM embeddings with structural features to inject geometry into sequence representations, improving downstream performance without learning at all-atom resolution. Representative methods include SaProt (Su et al., 2023b), FoldToken (Gao et al., 2024), ProSST (Li et al., 2024), and ESM3 (Hayes et al., 2024). Most hybrid models augment sequence tokens with backbone-level descriptors, and the learned tokens remain sequence-anchored. SLAE instead learns *structure and energetics-anchored residue tokens*, reducing sequence-only bias while increasing structure representation resolution.

### All-atom Protein Representation

All-atom protein generative models which simultaneously generate backbone and side chain coordinates can also have an all-atom representation of protein structure. Protpardelle (Chu et al., 2024) can be cast as a continuous normalizing flow to generate deterministic latent encodings of all-atom protein structures. A joint embedding space of sequence and all-atom structure was proposed in CHEAP (Lu et al., 2024), in which the embeddings reconstruct all-atom protein structures and recover sequence. However, interpolation between two conformations of the same protein sequence is not possible as identical sequence would map to the same CHEAP embedding. Representations can also be derived from protein structure prediction models such as AlphaFold3 (Abramson et al., 2024), but the information is distributed across layers and in both single and pairwise representations.

### Geometric GNNs for Atomistic Systems

Representing atomistic systems as geometric graphs is natural. While encoders for protein have been proposed using point cloud voxelization, graph convolution and hierarchical pooling (Hermosilla et al., 2021; Anand et al., 2022; Wang et al., 2023), they incur a considerable computational burden making them impractical for large-scale pretraining with previously proposed denoising objectives. Equivariant GNNs such as DimeNet (Gasteiger et al., 2022), NequIP (Batzner et al., 2022) and MACE (Batatia et al., 2023) excel at small-molecule property prediction and interatomic potentials. For scalability, many adopt low-order interactions with truncated neighborhoods, closely related to Atomic Cluster Expansion (ACE) formulations (Drautz, 2019). Works which extend atomistic modeling to proteins are emerging (Pengmei et al., 2024; Bojan et al., 2025), but existing approaches typically pretrain on small-molecule datasets, reuse features from pretrained potential models or are trained in a task-specific manner. There remains a gap in methods amenable to large-scale, all-atom pretraining on proteins. SLAE addresses this by *modeling two-body local interactions over cutoff graphs and pretrain a physics-informed autoencoder* that yields a general, task-agnostic latent space at protein scale: thousands of atoms per system compared to tens of atoms.

## 3 The SLAE Framework

We introduce the SLAE autoencoder and its end-to-end pretraining objectives (Fig. 1A). SLAE solves a deliberately difficult two-part problem: the geometric graph encoder projects interatomic interactions within each atom’s local neighborhood into compact residue tokens, while the decoder learns a global prior over how these local environments compose into coherent macromolecular structures. This residue-level bottleneck over all-atom inputs makes large-scale pretraining tractable and learns meaningful embeddings.

**Figure 1.**
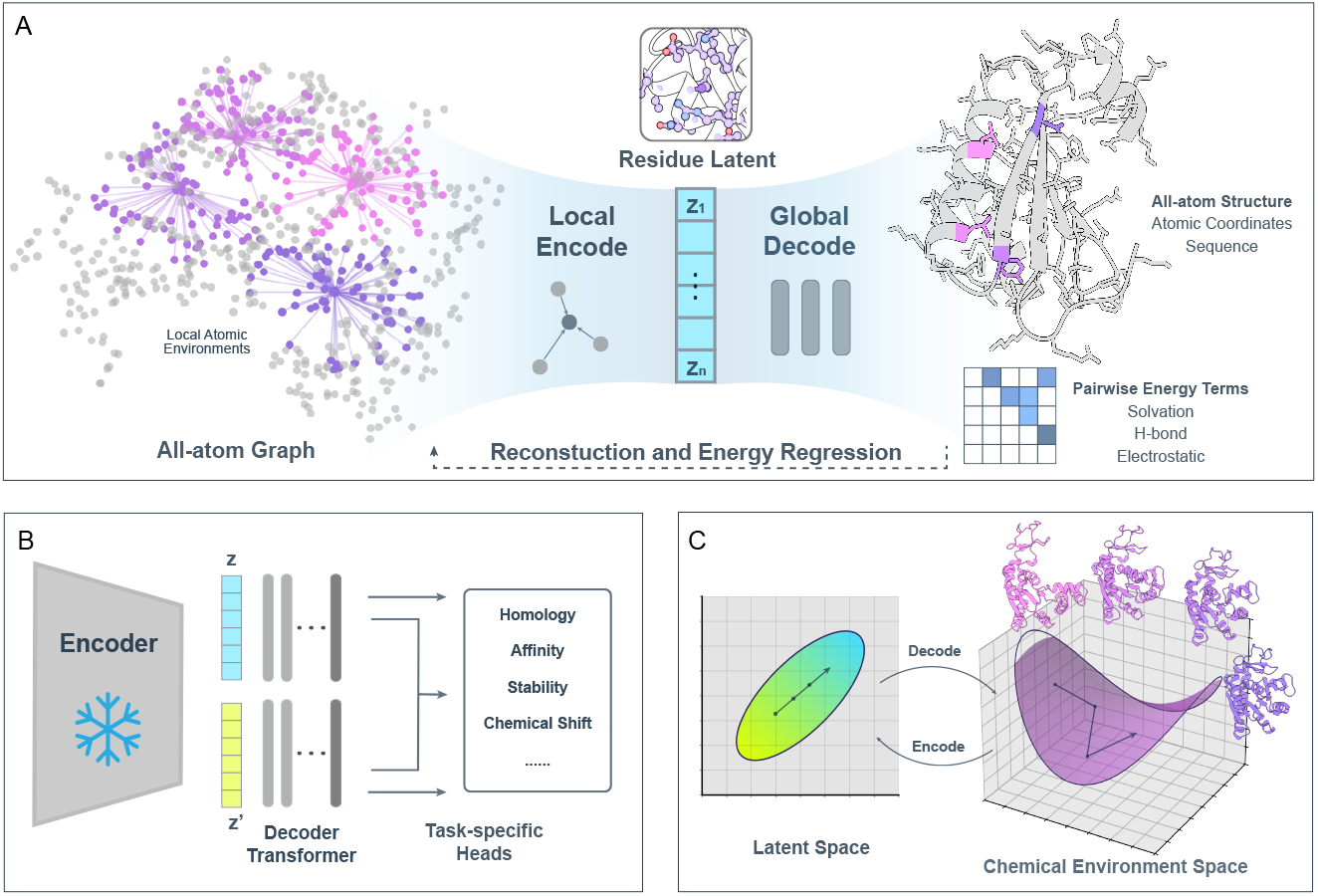
Overview of the SLAE framework. **A. Pretraining** A graph encoder maps local atomic neighborhoods to residue embeddings. Examples of atom connectivity shown as input to the encoder, with different colors for each residue. The transformer decoder connects pooled local features at residue level into the full-atom protein structure. The decoder also regresses to inter-residue energy score terms. **B. Transfer learning** The pretrained embeddings are fed to lightweight heads for diverse downstream tasks. **C. Latent geometry** Linear interpolations on latent space decode to physically coherent structures that follow changes on the underlying chemical-environment manifold.

### 3.1 Structure Representation

Given a protein structure, we construct a directed graph 𝒢 = (𝒱, *ℰ*), where:

#### Nodes

Each node *v*_*i*_ ∈ 𝒱 represents heavy atom *a*_*i*_. The node feature is a one-hot encoding of the atom’s chemical type.

#### Edges

For each pair of atoms *a*_*i*_, *a*_*j*_ with ∥***a***_*j*_ − ***a***_*i*_∥_2_ ≤ 8Å, we define a directed edge *e*_*j*→*i*_ ∈ *ℰ* with features 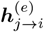 that is a concatenation of: (i) the scaler *interatomic distance* ∥***a***_*j*_ − ***a***_*i*_∥_2_ in terms of Bessel radial basis functions *ϕ*_*r*_(***a***_*i*_, ***a***_*j*_) and (ii) the unit vector *interatomic direction* projected onto spherical harmonics *Y*_*ℓm*_ *ϕ*_*a*_(***a***_*i*_, ***a***_*j*_).

#### Design Motivation

This representation is *minimal yet physically complete*: it encodes interatomic distances and orientations without relying on torsion angles, amino acids, or residue indices. As such, it enables generalization to arbitrary biomolecular complexes, which we leave for future work. Bond connectivity and hydrogen patterns are learned implicitly through the autoencoder objective detailed in Section 3.4.

### 3.2 Encoder

The encoder maps each atom’s local chemical environment into residue-level latent embeddings {***z***_1_, …, ***z***_*n*_}, ***z***_*i*_ ∈ ℝ^128^.

#### Equivariant Neighborhood Embedding

We employ a SE(3)-equivariant neural network, inspired by Musaelian et al. (2023), that operates on each heavy atom and its neighbors through learned edge embeddings. Each layer *L* maintains coupled latent spaces: a scalar space 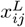 (invariant) and a tensor space 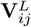 (equivariant). An equivariant tensor product incorporates interactions between the current equivariant state of the center–neighbor pair (*i, j*) and all other neighbors 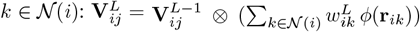, where *ϕ*(**r**_*ik*_) is a geometric embedding of the neighbor direction and 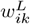 are learned weights derived from scalar features of edges (*i, k*). This can be viewed as a weighted projection of the atomic density around atom *i*, enabling equivariant interactions between the pair (*i, j*) and the environment of *i*.

Following the tensor product, scalar outputs are reintroduced into the scalar latent space with 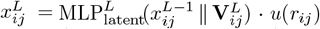, where *u*(*r*_*ij*_) is a smooth cutoff envelope. This step completes the coupling of scalar and equivariant latent spaces: scalars distilled from tensor products inject directional information back into 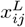, allowing the invariant channel to carry geometric cues that were previously only available to the equivariant representation.

#### Residue Environment Pooling

After the final layer, we obtain scalar pair features 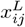. We first pool to atoms by mean-aggregating incoming edges, and then pool atom embeddings to residues: 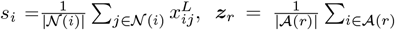. This yields compact residue-level representations while retaining strictly local chemical information.

#### Design Motivation

The encoder updates edge embeddings dynamically by incorporating information from neighboring edges. This paradigm originally developed for interatomic potentials in smallmolecule graphs naturally extends to large protein graphs. This allows SLAE to capture strictly local but physically meaningful chemical environments. Pooling representations to the residue level serves as an efficient and natural information bottleneck for protein structure.

### 3.3 Decoder

Having distilled each residue’s local chemistry and geometry into embeddings ***z*** ∈ ℝ^128^, the decoder assembles these local descriptors into a single, coherent macromolecule that respects long-range couplings.

#### Architecture

We first project each latent embedding to a model dimension of ℝ^1024^. On top of these expanded embeddings, we employ a Transformer architecture with global self-attention and Rotary Positional Embeddings (RoPE) (Su et al., 2023a) to capture long-range residue interactions with a stack of multi-head self-attention layers.

The Transformer outputs are passed into three parallel MLP heads for structure reconstruction, sequence recovery, and energy prediction:

1. Reconstructs the 3D coordinates of up to 37 heavy and side-chain atoms per residue 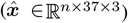.
2. Recovers the amino acid identity at each residue position (***ŝ*** ∈ ℝ^*n×*20^).
3. Approximates inter-residue physical interactions using Rosetta scores, including hydrogen bonding, electrostatics, and solvation energies 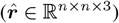.

#### Design Motivation

The decoder is designed to complement the encoder’s strictly local representation by modeling *global* dependencies across residues. Global self-attention allows residue embeddings to exchange information across the entire protein, enabling the reconstruction of coherent backbone and side-chain geometries. The addition of energy prediction task guides the decoder toward physically meaningful structures, ensuring that the latent space encodes not only geometric detail but also the energetic constraints that govern protein stability and interactions.

### 3.4 Pretraining

We pretrain SLAE end-to-end on full atomic structures with three complementary objectives:

1. **All-atom Structure Recovery** To obtain the predicted structure 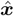, we mask the atom-37 template coordinates while providing the ground-truth residue identities to train the decoder to recover ground truth coordinates. We supervise this reconstruction with a combination of all-atom local distance difference test loss (SmoothLDDT) (Jumper et al., 2021) and frame-aligned point error (FAPE) (Anishchenko et al., 2024): 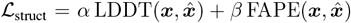, where ***x*** and 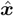 denote the ground-truth and predicted all-atom coordinates.
2. **Sequence Recovery** We additionally recover the residue sequence from the latent space: 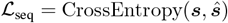, where ***s*** is the ground-truth amino-acid identity and ***ŝ*** are the predicted logits over 20 amino acid classes.
3. **Energy Prediction** To inject physically grounded supervision, we predict inter-residue energies approximated by Rosetta scores, including hydrogen bonding, electrostatics, and solvation: 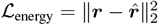, where ***r*** and 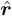 are ground-truth and predicted energy terms.

The combined loss integrates all three components:

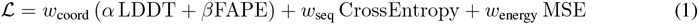

with weights *w*_struct_, *w*_seq_, *w*_energy_ ≥ 0 as tunable hyperparameters (Appendix B.1).

#### Implicit Latent Space Regularization

By jointly optimizing geometry, identity, and energetics, SLAE’s pretraining objective provides complementary constraints on the latent space: **(i)** Geometry losses depend smoothly on atomic coordinates, promoting continuous and physically plausible reconstructions. **(ii)** Sequence recovery encourages embeddings to encode amino acid identity, preserving biochemical interpretability and avoiding collapse. **(iii)** Energy prediction provides a physics-based signal, guiding embeddings toward inter-residue interactions such as hydrogen bonding, solvation, and electrostatics. These losses shape a latent manifold that maps cleanly onto valid, physically coherent protein conformations. The result is a structurally consistent, chemically informative, and energetically grounded representation without relying on explicit regularizers.

### 3.5 Results and ablations

We pretrain SLAE on a sequence-augmented CATH(Ingraham et al., 2019)-derived dataset (Lu et al., 2025b)(Appendix C). On the held-out test set with no family overlap, the autoencoder achieves 99.9% sequence recovery and all-atom RMSD of 1.1Å for structures shorter than 128 residues and 1.9Å across all lengths up to 512 residues.

We study the effect of model and pretraining design choices on pretraining performance (Table 6).

For encoder locality, we swept cutoff radii and find an 8 Å neighborhood yields the best results (Appendix E). For discretization, we compare end-to-end VQ (van den Oord et al., 2018) and LFQ (Yu et al., 2023) against post-hoc *k*NN codebooks built on frozen encoder embeddings. End-to-end quantization trades off sequence and structure accuracy, whereas reconstruction from post-hoc *k*NN-codebook quantized embeddings approaches continuous resolution as the codebook grows. Ablation experiments (Table 6, Appendix E) further highlight the importance of both the FAPE loss and Rosetta-derived energy supervision, confirming the effectiveness of our multitask pretraining framework. These results validate the design choices and permit downstream evaluation on a faithful representation of protein structures.

## 4 Downstream Tasks

We next demonstrate that SLAE embeddings pretrained on all-atom reconstruction and energetics objectives transfer effectively to diverse downstream tasks(Figure 1B). Across all four benchmarks spanning complementary biological scales, SLAE achieves better or on-par performance with state-of-the-art methods, underscoring the generality and flexibility of the SLAE framework.

### Fold Classification

Protein fold classification is a cornerstone of structural biology, linking structure to evolutionary relationships and functional annotation. Using the SCOPe 1.75 dataset Fox et al. (2014) and following Hou et al. (2018), we evaluate generalization under three test sets: Family, Superfamily, and Fold. An MLP is trained on pooled residue embeddings. SLAE achieves on-par or superior accuracy compared to prior state-of-the-art models across all splits (Table 2), demonstrating that global fold information can be recovered even from strictly local all-atom embeddings.

**Table 1.** Reconstruction performance of SLAE and ablations. We report sequence recovery accuracy (%) and reconstruction RMSD (Å) on test structures. All further experiments use the highlighted best SLAE model.

| Graph Radius(Å) | Discretization Method | Codebook Size | Training Obj. | Seq. Acc. (%) | RMSD < 128 res (Å) | RMSD < 512 res (Å) |
| --- | --- | --- | --- | --- | --- | --- |
| 8 | LFQ | 32768 | all | 75.2 | 2.50 | 3.74 |
| 8 | kNN | 4096 | all | 97.5 | 2.96 | 4.03 |
| 8 | kNN | 32768 | all | 99.4 | 1.60 | 2.31 |
| 8 | – | – | w/o. FAPE | 97.2 | 3.89 | 5.22 |
| 8 | – | – | w/o. Energy | 98.0 | 3.26 | 5.17 |
| 6 | – | – | all | 99.9 | 1.24 | 2.55 |
| **8** | **–** | **–** | **all** | **99.9** | **1.12** | **1.92** |

**Table 2.** Fold classification accuracy (%) on SCOPe 1.75 under three test splits.

| Method | Fold (%) | Superfamily (%) | Family (%) |
| --- | --- | --- | --- |
| GVP-GNN(Jing et al., 2021) | 16.0 | 22.5 | 83.8 |
| IEConv(Hermosilla et al., 2021) | 45.0 | 69.7 | 98.9 |
| GearNet-Edge-IEConv (Zhang et al., 2023) | 48.3 | 70.3 | 99.5 |
| ProNet-SCHull (Wang et al., 2024) | **56.1** | 74.6 | **99.4** |
| SLAE-finetuned | 55.1 | **77.1** | 99.1 |

### Protein-Protein Binding Affinity Prediction

Protein-protein interactions underlie nearly all cellular processes, and accurate prediction of binding affinity is critical for understanding signaling pathways, complex assembly, and therapeutic design. We evaluate SLAE on the PPB-Affinity dataset (Liu et al., 2024), a recently curated large-scale benchmark that aggregates 12,062 experimental binding ΔΔ*G* values from multiple sources and aligns them with high-quality structural complexes.

Complex structures are embedded chain-wise and interface-wise with the SLAE encoder, and pooled residue embeddings are passed into an MLP for regression. In 5-fold cross-validation, SLAE achieves lower RMSE and higher Pearson correlation than PLM-based baselines (Table 3). Despite being pretrained only on single-chain data, SLAE generalizes seamlessly to multi-chain contexts, thanks to its atomistic representation that does not rely on residue or chain indices.

**Table 3.** Protein-protein binding affinity prediction on the PPB-Affinity dataset.

| Method | RMSE (kcal/mol) | Pearson Correlation |
| --- | --- | --- |
| PPB-Affinity Baseline (Liu et al., 2024) | 2.08 | 0.70 |
| PPLM-Affinity (Liu et al., 2025) | 1.89 | 0.76 |
| SLAE-finetuned (w/o. interface) | 2.01 | 0.73 |
| SLAE-finetuned (with interface) | **1.86** | **0.77** |

### Single-Point Mutation Thermostability Prediction

Protein stability is fundamental to function, and predicting the impact of point mutations on thermostability (ΔΔ*G*) is a central challenge for protein engineering, drug resistance modeling, and disease variant interpretation. We benchmark SLAE on the Megascale mutation dataset (Tsuboyama et al., 2023), filtered according to ThermoMPNN protocol with 272,712 mutations across 298 proteins Dieckhaus et al. (2024).

Pairs of wild-type and mutant structures are embedded with residue-level differences extracted at the mutation site. An MLP head predicts ΔΔ*G*. SLAE achieves 0.68 RMSE and 0.76 Pearson correlation (Table 4) on the test set, outperforming prior methods. Ablation experiments show that removing mutation-site differencing degrades performance, highlighting the importance of local residue environment modeling for physical property prediction in the SLAE framework.

**Table 4.** Single-point mutation thermostability prediction on the Megascale dataset test split.

| Method | RMSE (kcal/mol) | Pearson Correlation |
| --- | --- | --- |
| Rosetta (Pancotti et al., 2022) | 5.18 | 0.53 |
| RaSP (Blaabjerg et al., 2023) | 1.08 | 0.71 |
| ThermoMPNN (Dieckhaus et al., 2024) | 0.71 | 0.75 |
| SLAE-finetuned (w/o. mutated site) | 0.73 | 0.70 |
| SLAE-finetuned (with mutated site) | **0.68** | **0.76** |

### Chemical Shift Prediction

NMR chemical shifts are among the most direct experimental probes of local atomic environments, among them the backbone nitrogen are notoriously difficult to predict accurately due to its large variance and contributions from ring currents, electrostatics, and subtle side-chain conformations. We benchmark on stringently filtered BMRB (Hoch et al., 2023) which contains 2,532 training and 594 validation chemical shift records and their corresponding Alphafold2 predicted structures. PLM-CS framework is adopted as baseline model architecture, which trains a lightweight predictor on top of pretrained representations (Zhu et al., 2025).

We report validation set performance of finetuned SLAE along with PLM-CS results using multiple protein residue embeddings, including ESM2, AlphaFold2, ProSST and SLAE. ^1^ Finetuned SLAE achieves the lowest RMSE and highest correlation, substantially outperforming retrained PLM-CS baselines (Table 5). This demonstrates that SLAE embeddings capture fine-grained atomistic features essential for NMR observables.

**Table 5.** Backbone nitrogen chemical shift prediction on BMRB.

| Method | RMSE (ppm) | Pearson Correlation |
| --- | --- | --- |
| PLMCS-AF2 | 2.94 | 0.82 |
| PLMCS-ESM2 | 2.74 | 0.84 |
| PLMCS-ProSST | 2.53 | 0.87 |
| PLMCS-SLAE | 2.53 | 0.87 |
| SLAE-finetuned | **1.88** | **0.93** |

## 5 Interpreting the Latent Space

SLAE’s downstream performance stems from a structured, interpretable latent space. We show that residue embeddings are organized along biochemically meaningful axes, are sensitive to local environment changes, and admit linear paths that decode to geometrically coherent structures(Figure 1C).

### 5.1 Embedding Variability Reflects Chemical Environment Change

To probe what SLAE embeddings captures at the residue level, we analyze how they organize across local chemical environments. Dimensionality reduction of *k*NN centroids from CATH (Section 3.5, Appendix E) shows that residue latents cluster by side chain chemistry and broader structural context. The latent space also stratifies along gradients of solvent accessibility and separates by secondary structure, with helices, sheets, and coils occupying distinct submanifolds (Figure 3, App. Fig 6 and 7). This indicates that SLAE representation is sensitive to both chemical identity and structural environment.

**Figure 2.**
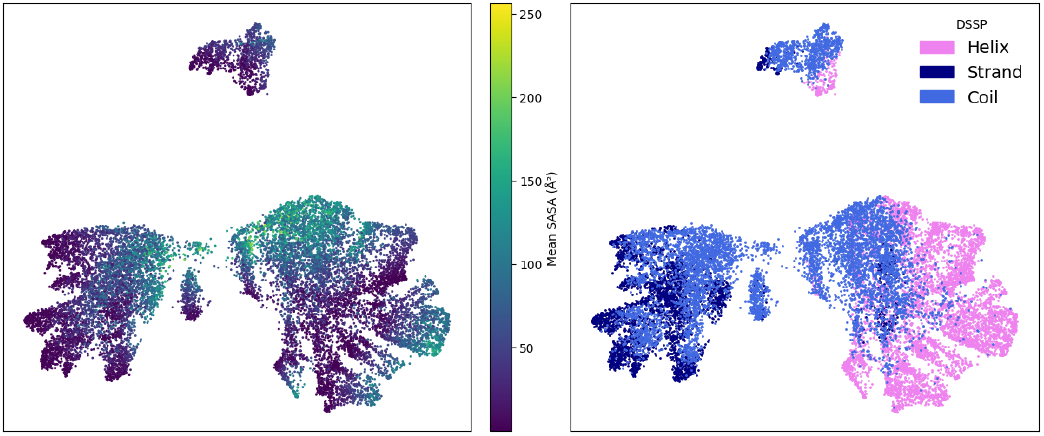
SLAE latent organization. UMAP visualization of *k*NN centroids shows clustering by solvent accessibility (left) and secondary structure (right).

**Figure 3.**
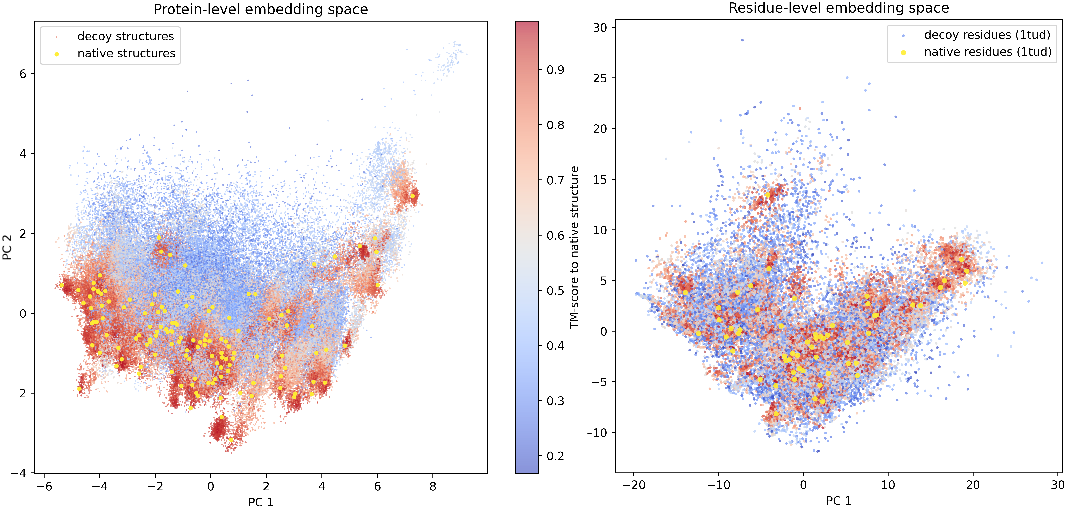
SLAE embedding comparison between native and decoy structures. (native in yellow, decoys colored by TM-score to their native; warmer = more native-like) **Left** protein-level PCA. Each point is a protein. **Right** residue-level PCA for 1TUD and its decoys. Decoy residues are colored by their parent decoy’s TM-score. In both panels, SLAE embeddings organize along gradients of nativeness, revealing coherent neighborhoods that align with structural quality.

We then quantify this sensitivity using the mdCATH dataset (Mirarchi et al., 2024). Across 5,398 proteins, per-residue latent displacement between conformers correlates with physical measures of environment variability: changes in contact maps and solvent exposure explain over half of the variance in embedding similarity (*R*^2^ = 0.55, *ρ* ≈ 0.74; Appendix E). Thus, SLAE embeddings consistently track how residues respond to burial, packing, and secondary-structure transitions.

### 5.2 Discriminative power over Native-Decoy Residue environments

We show that SLAE residue latent capture local environments contain signal that zero-shot distinguishes native structures from decoys and provide a practical embedding space for evaluating backbone–sequence co-design.

On the Rosetta decoy dataset (Park et al., 2016) containing 133 native protein structures with thousands of decoys each, native–decoy cosine margin is 0.136 across residues. We further fit a leaveprotein-out logistic regression by training on all proteins except one and tested on the held-out protein’s residues and report AUROC = 0.659 (Appendix E), indicating a moderate, generalizable linear signal at the residue level.

Motivated by this discriminative signal, we use the SLAE embedding space to quantify the distributional coverage of generative models, extending prior metrics (Lu et al., 2025a) to all-atom resolution and residue granularity. As a proof of concept, we compute per-residue type Fréchet Protein Distance (FPD) between SLAE embeddings of the generated structures and the native CATH distribution for models such as Chroma (Ingraham et al., 2023), Protpardelle-1c (Lu et al., 2025b) and La-Proteina (Geffner et al., 2025). The FPD metrics reveal subtle differences in the coverage of local amino acid environments by different generative models (Appendix E.3, App.Fig. 8). For example, biased sampling is evident in La-Proteina samples for serine, threonine, and valine relative to Protpardelle-1c and Chroma. Using SLAE embeddings provides a more sensitive view on coverage of all-atom local environments which are ignored in backbone-based metrics and which may be averaged out on the global protein fold level as in previous assessments of generative model coverage of protein structures

### 5.3 Smooth Latent Interpolation Captures Conformational Transitions

Latent space smoothness is relevant for evaluating whether a representation supports continuous sampling of protein conformations. Unlike variational autoencoders that encourage smoothness via KL regularization to a simple prior, the SLAE autoencoder relies solely on physics-augmented pretraining objectives. We examine the smoothness of SLAE latent by linear interpolation between two conformation states ***Z***^(*A*)^ and ***Z***^(*B*)^. For each residue *i* and interpolation scale *t* ∈ [0, 1], the interpolated residue embeddings are given by 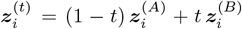. The interpolated set 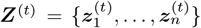 is then decoded into an all-atom structure with the pretrained SLAE decoder (Figure 1C).

For two proteins with known conformational changes, adenylate kinase (AdK) and KaiB, we linearly interpolate between the SLAE embedding of the two experimentally determined states(AdK: 1AKE, 4AKE; KaiB: 2QKE, 5JYT). We sample intermediate structures from 50 evenly spaced values of *t* and align their backbone coordinates to frames in MD simulation of the transitions (Seyler et al., 2015; Zhang et al., 2024). For AdK, the interpolated structures closely track the MD intermediates, as evidenced by smooth trajectories with low RMSD (Figure 4), and they agree better than interpolations from the generative model (App. Fig. 10). Notably, these interpolations are *unguided by any energy function or model likelihood*; they arise solely from linear paths in SLAE latent space anchored in pretraining with physics-based task. KaiB shows higher RMSD between steps 20 and 30 (Figure 4). Closer examination of the interpolated structures (App. Fig 9) reveals disagreement in the C-terminus, which is known to unfold during transition (Wayment-Steele et al., 2023). This degradation is expected as SLAE is pretrained on folded structures and thus treats unfolded segments as out-of-distribution, where local environment cues under-constrain reconstruction.

**Figure 4.**
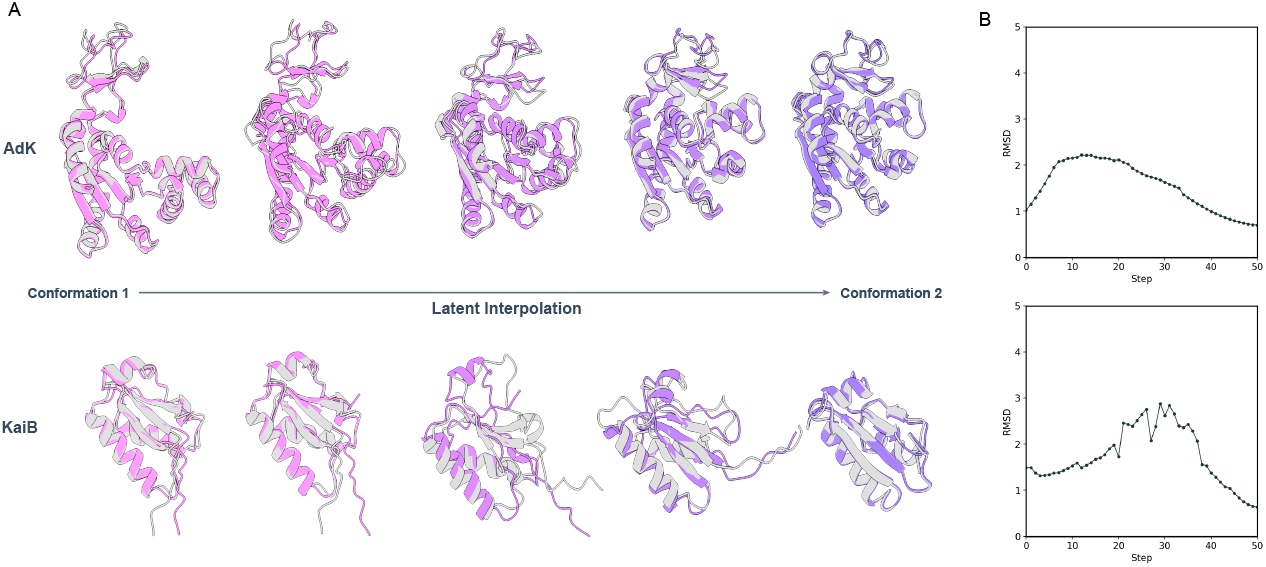
Latent space interpolation between two conformations A. Structures sampled by linear interpolation (purple) overlaid with MD simulation frames (grey) **B.** Alignment RMSD to MD simulation trajectories

Within the folded structure regime, SLAE’s latent space is sufficiently regular that simple linear paths often decode to geometrically coherent intermediates aligned with MD trajectories. These results support the view that SLAE embeddings approximate a continuous, chemically grounded manifold of protein structures. The latent space reflects local environmental variation while accommodating large-scale transitions, make it useful for downstream analysis and generative applications.

## 6 Conclusion

We introduced SLAE, a framework tailored to learning general-purpose representations of proteins at all-atom resolution. SLAE applies a strictly local graph neural network over atomic environments, using computationally simple layers to perform expressive geometric reasoning on atom-type and interatomic distance features. Pretraining is driven by a novel objective that combines full atomic coordinate reconstruction with energy score regression, yielding embeddings that are structurally faithful, chemically grounded, and energetically informed.

## Acknowledgments

The computing for this project was performed on the Sherlock cluster. We would like to thank Stanford University and the Stanford Research Computing Center for providing computational available resources and support that contributed to these research results. Y.C and T.L. are supported by Stanford Graduate Fellowship. C.Z. and H.K.W.-S. acknowledge financial support from the University of Wisconsin-Madison Office of the Vice Chancellor for Research, with funding from the Wisconsin Alumni Research Foundation. This project is supported by NIH (R01GM147893 to P.-S.H.), Merck Research Laboratories (MRL) Scientific Engagement and Emerging Discovery Science (SEEDS) Program, and Stanford Medicine Catalyst. The views and conclusions contained in this document are those of the authors and should not be interpreted as representing the official policies, either expressed or implied, of the U.S. Government.

## A Model

### A.1 Autoencoder pseudocode

The end-to-end SLAE autoencoder can be summarized as follows:

#### Algorithm 1

SLAE Autoencoder. ***h***^(*e*)^: edge features, ℰ : SE(3)-equivariant update, 𝒫: pooling, 𝒟_Tr_: Transformer decoder. Outputs: 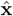 (coordinates), ***ŝ*** (sequence), 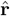 (energies).

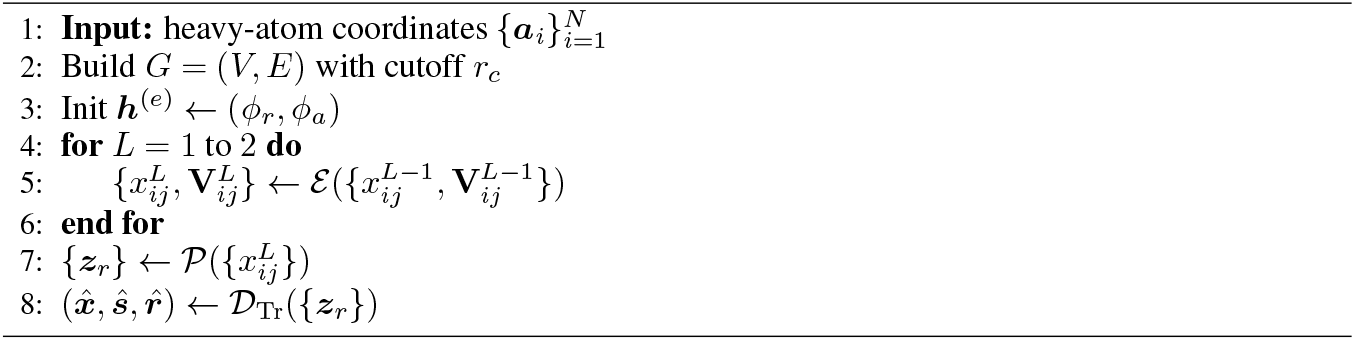

### A.2 Encoder Architecture

**Notation** Let ***a***_*i*_ ∈ ℝ^3^ be the coordinate of atom 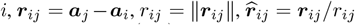. The neighbor set of *i* is (*i*) ={ *j* | *r*_ij_≤ *r*_*c*_}. Each directed edge (*i, j*) maintains invariant scalars 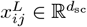 and equivariant tensors 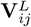.

#### Two-body initialization

Edge features are initialized with radial and angular bases:

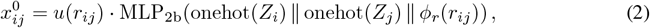

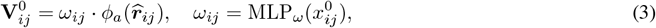

where *ϕ*_*r*_ are Bessel radial basis functions, *ϕ*_*a*_ are angular embeddings (e.g., spherical harmonics), and *u*(*r*_*ij*_) is a smooth cutoff envelope.

#### Tensor product update

At layer *L*, equivariant features of edge (*i, j*) interact with the embedded environment of atom *i*:

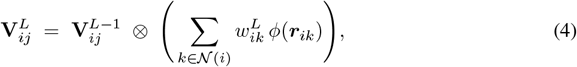

where *ϕ*(***r***_*ik*_) encodes neighbor geometry and 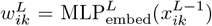 are learned weights. This corresponds to a weighted projection of the atomic density around atom *i*.

#### Latent scalar update

Scalar channels are updated with tensor product scalars:

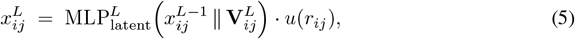

injecting geometric information from 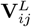 back into 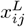.

#### Hierarchical pooling

Final edge scalars are aggregated:

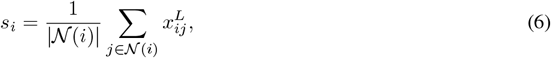

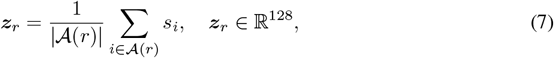

producing residue-level embeddings {***z***_*r*_}.

### A.3 Decoder architecture

#### Transformer backbone

We employ a standard pre-norm Transformer encoder with Rotary Positional Embeddings (RoPE) with *L*_Tr_=8 layers, *h*=16 heads, model width *d*_model_=1024. Each layer consists of:

- Multi-head self-attention with RoPE (pre-norm): MHA_RoPE_(LayerNorm(·)).
- Residual connection.
- Feed-forward network with hidden dimension *d*_ff_ and SwiGLU, applied as FFN(LayerNorm(·)).
- No dropout.

Formally:

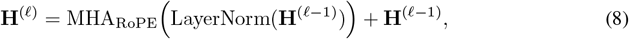

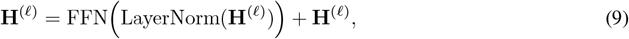

for 𝓁 = 1, …, L_Tr_, with H^(0)^ = [z_1_, …, z_n_].

#### Prediction heads

From final hidden states 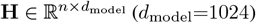, we apply three parallel heads:

i. **3D coordinates (linear head)** LayerNorm + Linear maps per-residue embeddings to all Atom37 coordinates:

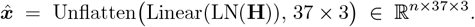

(Atoms 1–4 are N, C_*α*_, C, and O; atoms 5–37 are side chain. Masking is applied via the Atom37 mask.)
ii. **Sequence logits on valid tokens** An MLP head operates only on valid tokens (mask-compacted), then is re-padded for loss:

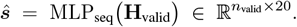 **Pairwise energies** A pairwise feature head first down-projects **H**, lifts to 2D by pairwise product/difference, applies a small MLP, then per-type linear heads with magnitude clamp to 1*e* − 3:

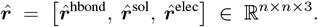

### A.4 Task-specific heads

#### Trainable decoder backbone

We expose a lightweight wrapper over the `DecoderBackbone` to enable fine-tuning the last *N* Transformer blocks while freezing the rest. Take the single site mutation stability task as an example, we document the layout of downstream task-specific finetuning here.

#### Contrastive and site-aware head

A Siamese head takes two or more structure embeddings (e.g., wild-type and mutant), runs them through the shared `DecoderBackbone`, and regresses a scalar target (e.g., ΔΔ*G*). Beyond global contrastive pooling, it can extract *site-specific* residue representations, enabling residue-level tasks.

#### Backbone embeddings

Given masked inputs (**X**^wt^, **M**^wt^) and (**X**^mut^, **M**^mut^),

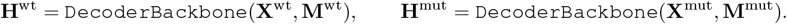

#### Mask-aware pooling and site features

Let the mean-pooling operator be

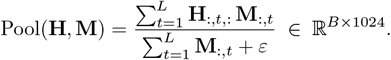

We form global embeddings **z**^wt^ = Pool(**H**^wt^, **M**^wt^) and **z**^mut^ = Pool(**H**^mut^, **M**^mut^). Given mutation indices ***ι*** ∈ {1, …, *L*}^*B*^, we also extract site embeddings

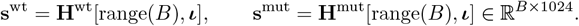

#### Contrastive feature and MLP regressor

We concatenate global and site representations together with their difference:

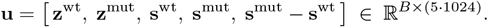

A small MLP head predicts a scalar per pair:

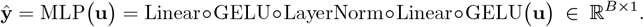

#### General usage

The same interface supports other pairwise or single-input tasks by (i) choosing one or multiple passes through `DecoderBackbone`, (ii) selecting global vs. site-wise features, and (iii) swapping the final MLP for the appropriate output dimensionality/loss. For atom-level tasks, the `DecoderBackbone` can be reinitialized with the smaller attention window.

## B Training

### B.1 Losses

#### All-atom FAPE (Frame-Aligned Point Error)

All-atom FAPE is computed by aligning the predicted and reference structures on every triplet of bonded atoms (*i, j, k*) (with the exception of symmetric side chain atoms) and then measuring per-atom positional deviations between the aligned structures. For each frame *f* (*i, j, k*) (with *j* as the origin), define an orthonormal basis for predicted/true coordinates via a deterministic map Φ : (ℝ^3^)^3^ → SO(3):

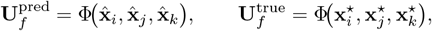

where Φ constructs column vectors from the two edge directions at *j*,

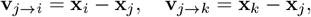

then

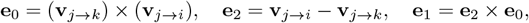

and column-normalizes [**e**_0_, **e**_1_, **e**_2_] to obtain a right-handed 3*×*3 matrix.

For any atom *a* in the same protein as *f*, rotate origin-subtracted positions into the local frames:

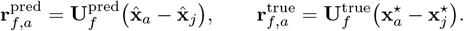

Define 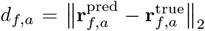, clamped at *c* = 10 A as 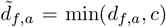, and apply a Huber penalty with *δ* = 1.0:

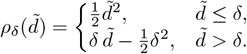

We average first over frames and then over atoms, yielding an atom-weighted mean:

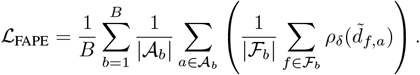

#### All-atom smooth LDDT

We use a differentiable, all-atom version of LDDT that compares pair-wise distances within a cutoff. Let 𝒫 = {(*i, a*), (*j, b*)} be all heavy-atom pairs with 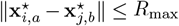 and not in the same residue. Define ground-truth and predicted distances 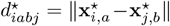 and 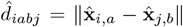, and the absolute error 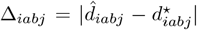. Using standard lDDT thresholds *τ*∈ { 0.5, 1.0, 2.0, 4.0} Å with smooth indicators *s*_*τ*_ (Δ) = *σ*(*α*(*τ* − Δ)) (sigmoid, *α* controls sharpness),

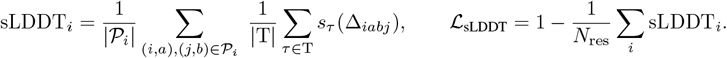

#### Mean-squared error (MSE)

Used for continuous targets regression:

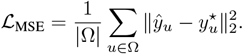

#### Huber loss

Used for continuous targets regression with *δ* = 1.35:

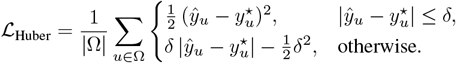

### B.2 Training specifics

The autoencoder is trained on a single NVIDIA A100 or H100 GPU using batch size 16. For pretraining we use *w*_coord_ = *w*_seq_ = *w*_energy_ = 1 and *α* = 10, *β* = 1 for the loss ℒ = *w*_coord_ (*α* LDDT + *β*FAPE) + *w*_seq_ CrossEntropy + *w*_energy_ MSE. We train for 30 epochs with early stopping on validation loss not decreasing after 5 epochs. The learning rate schedule is linear warmup for 1,000 steps followed by cosine decay. Optimization uses AdamW with maximum learning rate 1 *×* 10^−4^ and standard *β*_1_=0.9, *β*_2_=0.999 (weight decay as in AdamW defaults). Unless noted otherwise, downstream task-specific fine-tuning uses the same batch size and maximum learning rate 1 *×* 10^−5^.

## C Datasets

### Pretraining Structure

We train SLAE on an sequence-augmented CATH set (Lu et al., 2025b) by redesigning each domain with 32 ProteinMPNN sequences and predicting structures with ESMFold; we retain only high-confidence, self-consistent structure models (*pLDDT* ≥ 80, *scRMSD* ≤ 2.0*Å*), yielding 337936 structures, with 271 test structures from holdout CATH domains. We evaluate SLAE latent space on protein conformational ensembles sampled from the dataset of molecular dynamics (MD) simulations mdCATH (Mirarchi et al., 2024). We subsample 32 frames per protein across MD trajectory ensembles for each of the 5398 structures.

### Pretraining Rosetta Score

We use PyRosetta to compute residue pairwise energy scores for all pretraining structures under its default full-atom energy terms. For each pair of residues we compute (1) *fa sol*: Lazaridis-Karplus solvation energy (2) *fa elec*: Coulombic electrostatic potential with a distance-dependent dielectric (3) *hbond*: Sum of all hydrogen bonding terms for backbone and sidechain.

### Fold Classification

We obtain the dataset from Hermosilla et al. (2021), which consolidated 16,712 proteins with 1195 different folds from the SCOPe 1.75 database (Fox et al., 2014). Three test sets are used: (1) Family, which allows proteins from the same family to appear in both training and test; (2) Superfamily, which excludes proteins sharing family membership with the training set; and (3) Fold, which further excludes proteins from the same superfamily as those in training. All structures are obtained from the SCOPe 1.75 archive.

### Stability

We obtain the dataset curated by Dieckhaus et al. (2024) on Tsuboyama et al. (2023), composed of 272,712 single point mutations and their experimental ΔΔ*G*. The proteins were clustered using MMseqs2 with sequence identity cutoff of 25% to yield 239 training, 31 validation and 29 validation proteins. For wild type sequences we predict their structures with AlphaFold2. For all mutated structures we model the mutation with PyRosetta and relax within 8Å radius to obtain training structures.

### Binding Affinity

We use the PPB-AFfinity (Liu et al., 2024) which integrates experimental protein-protein binding affinity data from several source databases: SKEMPI v2.0, SAbDab, PDB-bind v2020, Affinity Benchmark v5.5, and ATLAS. This dataset contains 12062 unique binding complexes consisting of 3032 unique PDB codes and point mutations. We use the structures curated in the dataset and define interface residues as those within 5Å distance from other atoms of the neighboring chains. For all mutations we mutate the sidechain with PyRosetta and relax within 8Å radius to obtain training structures.

### NMR Chemical Shift

We retrieve the BMRB totaling 17,028 entries (2025-07-02) (Hoch et al., 2023). The entries were filtered and processed based on NMR experiment type, backbone chemical shift coverage, sequence consistency, basic experimental condition boundary plus any other routine re-referencing requirements. 3623 entries were retained and split into 2532 training and 594 validation entries at a 50% pairwise sequence-identity threshold after filtering entries without any nitrogen chemical shifts. Alphafold2 was used to generate all structures used in training.

### MD Simulation

For adenylate kinase (AdK), we use conformational ensembles generated using the Framework Rigidity Optimized Dynamics Algorithm (FRODA), yielding 200 trajectories (Seyler et al., 2015). For KaiB, we use the temperature-dependent fold-switching simulation from Zhang et al. (2024), subsampling every 10 frames out of the 4 successful fold-switching trajectories from the fold-switched state to ground state.

### Rosetta Decoy

To assess local residue environment embeddings distribution between native and decoy structure, we use structure dataset by Park et al. (2016), where each of the 133 native structures are accompanied with large numbers (≥ 1000 cluster centers) of alternative conformations (decoys).

## D Metrics

### Structure comparison

We report RMSD after optimal Kabsch rigid alignment for C*α*, backbone and all-atom. Given reference **X**^⋆^ ∈ ℝ^*n×*3^ and prediction 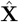, align 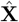 to **X**^⋆^ then compute

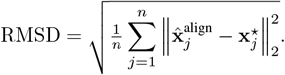

### Numeric regression

Given targets 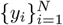 and predictions 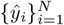, we report

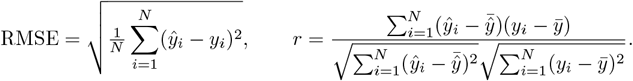

### Distribution comparison

We compute Fréchet Protein Distance (FPD) following Lu et al. (2025a). Given *N* data points from a reference distribution *p*_data_(**x**), here the sequence-augmented CATH dataset, and *M* samples from a generative model *p*_sample_(**x**), we computed per-residue SLAE embeddings 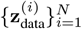 and 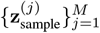 and then compute

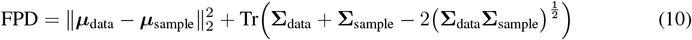

where ***µ***_data_ and ***µ***_sample_ are the means of the reference embeddings and the sample embeddings respectively, and **Σ**_data_ and **Σ**_sample_ are the covariance matrices of the reference embeddings and the sample embeddings respectively. We compute FPD using a smaller subset of 2000 samples as SHAPES showed that this is sufficient for an accurate FPD estimate Lu et al. (2025a).

## E Additional Experiments and Results

### E.1 Pretraining

We report in Table 6 additional results on the pretraining performance of the SLAE autoencoder. We note that encoders with 10Å graph radius cutoff is infeasible to train with a single GPU due to the number of edges.

**Table 6.**
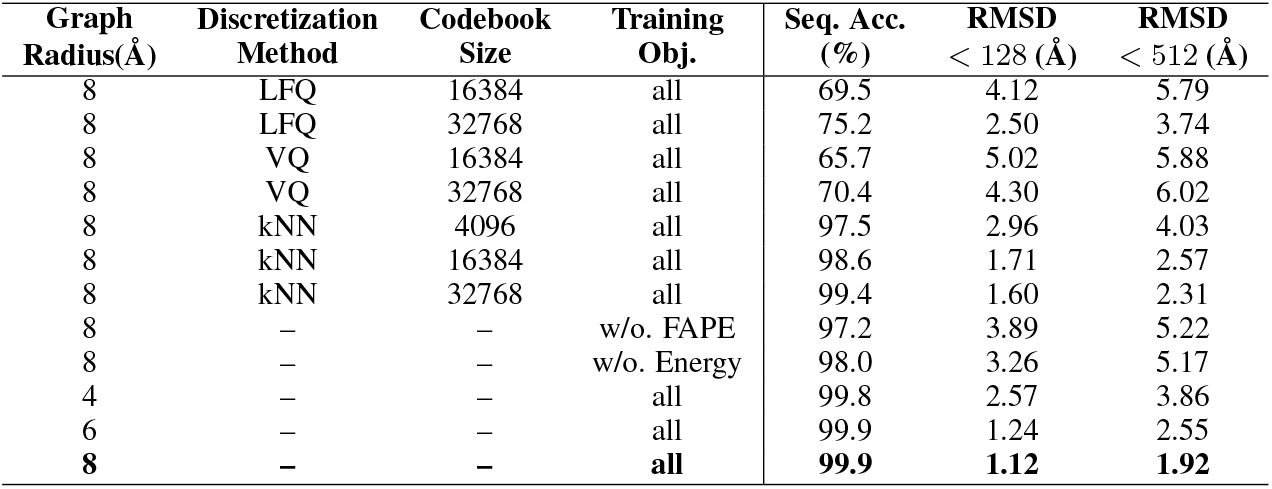
Complete results of SLAE autoencoder ablation experiments.

**Table 7.** Residue-wise clustering statistics: number of centroids that each residue type dominates, mean intra-cluster distance (*±* standard deviation), and ratio relative to the global mean.

| Residue | # Centroids | Mean intra-cluster distance | Ratio to global distance |
| --- | --- | --- | --- |
| | | mean ± std | |
| A | 1601 | 19.95 ± 6.50 | [1.20] |
| C | 215 | 17.67 ± 5.72 | [1.06] |
| D | 962 | 13.43 ± 7.47 | [0.81] |
| E | 1076 | 11.58 ± 3.97 | [0.70] |
| F | 641 | 11.44 ± 3.53 | [0.69] |
| G | 1192 | 18.38 ± 6.21 | [1.11] |
| H | 387 | 11.56 ± 3.53 | [0.70] |
| I | 899 | 14.33 ± 4.60 | [0.86] |
| K | 947 | 11.48 ± 3.76 | [0.69] |
| L | 1565 | 14.26 ± 4.63 | [0.86] |
| M | 272 | 13.69 ± 4.33 | [0.82] |
| N | 729 | 13.28 ± 4.16 | [0.80] |
| P | 737 | 13.83 ± 4.49 | [0.83] |
| Q | 492 | 11.48 ± 3.89 | [0.69] |
| R | 720 | 9.95 ± 3.14 | [0.60] |
| S | 1032 | 17.31 ± 5.58 | [1.04] |
| T | 920 | 15.81 ± 4.97 | [0.95] |
| V | 1253 | 15.96 ± 5.13 | [0.96] |
| W | 202 | 8.86 ± 2.76 | [0.53] |
| Y | 542 | 10.49 ± 3.16 | [0.63] |

### E.2 Latent space characterization

#### E.2.1 KNN clustering

We examine the CATH-kNN-quantized latent space, the k-means codebook of k = 16,384 centroids. We assign each centroid the majority amino-acid identity among its members; the commitment loss is the L2 distance from an embedding to its assigned centroid. The commitment loss histogram is tightly concentrated around 3–5 L2 units (Figure 5), which is modest relative to the embedding norm (15 *±* 4), indicating that quantization preserves most geometric signal.

**Figure 5.**
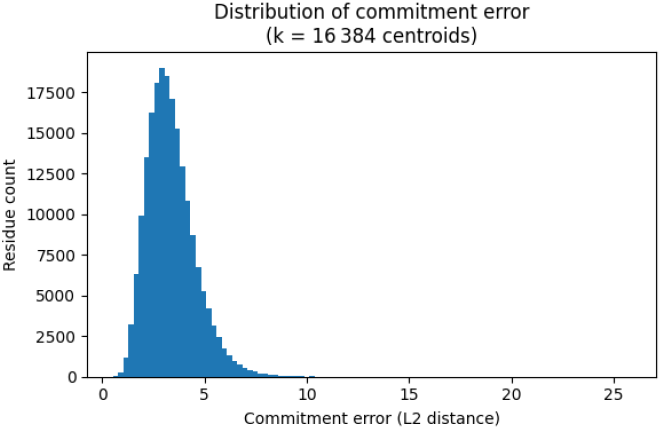
Commitment loss distribution during post-hoc quantization.

We observe clear residue type mixing in the clusters. Although many centroids are quite pure (median majority fraction 0.96), the distribution is broad (mean 0.89 *±* 0.15; entropy mean 0.52), with a substantial tail of mixed-composition clusters (10th-percentile majority 0.67). Along with the modest commitment error, this suggests that the observed mixing reflects genuinely overlapping local chemistries. Consistently, residue-conditioned intra-cluster distances show that some types form diffuse, mixed neighborhoods (A, G, S, C with ratios ≥ 1), while others are tighter and more typespecific (W, Y, R with ratios ≤ 1). These observations suggest that the kNN partitioning of residue embedding space yields chemically meaningful clusters but does not enforce one-residue exclusivity and captures real cross-type similarity in local environments.

#### E.2.2 Residue embedding visualization

We project the 16,384-entry codebook (centroid) embeddings into three dimensions using UMAP and analyze how local chemical environments are organized in this latent space (Figs. 6–7). Each CATH residue is assigned to its nearest codebook entry, and for every centroid we aggregate properties across its assigned residues. We compute the mean SASA and the majority secondary-structure label. This yields a coarse-grained landscape in which centroids arrange along solvent-exposure gradients and segregate by secondary-structure preferences.

**Figure 6.**
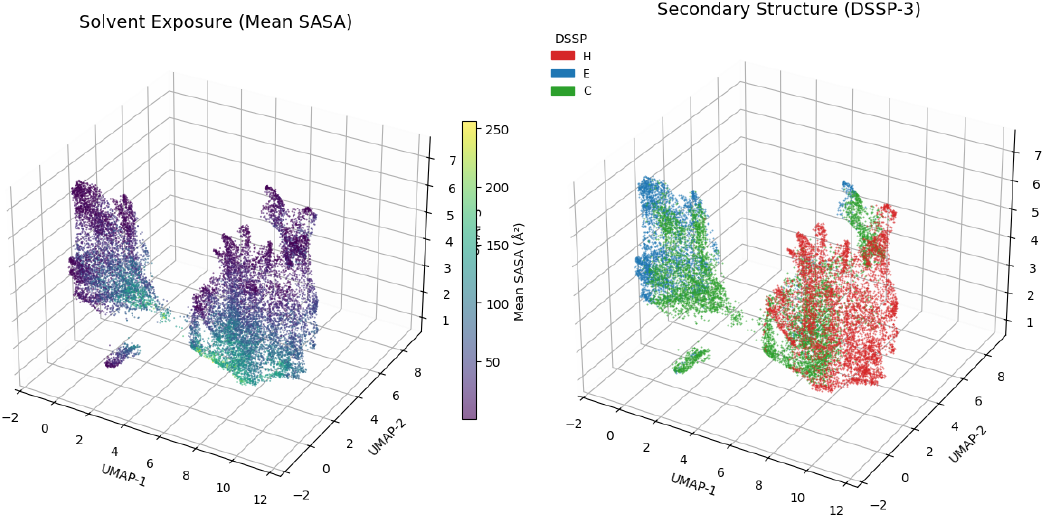
3D UMAP projection of CATH residue embeddings colored by solvent accessibility and secondary structure.

**Figure 7.**
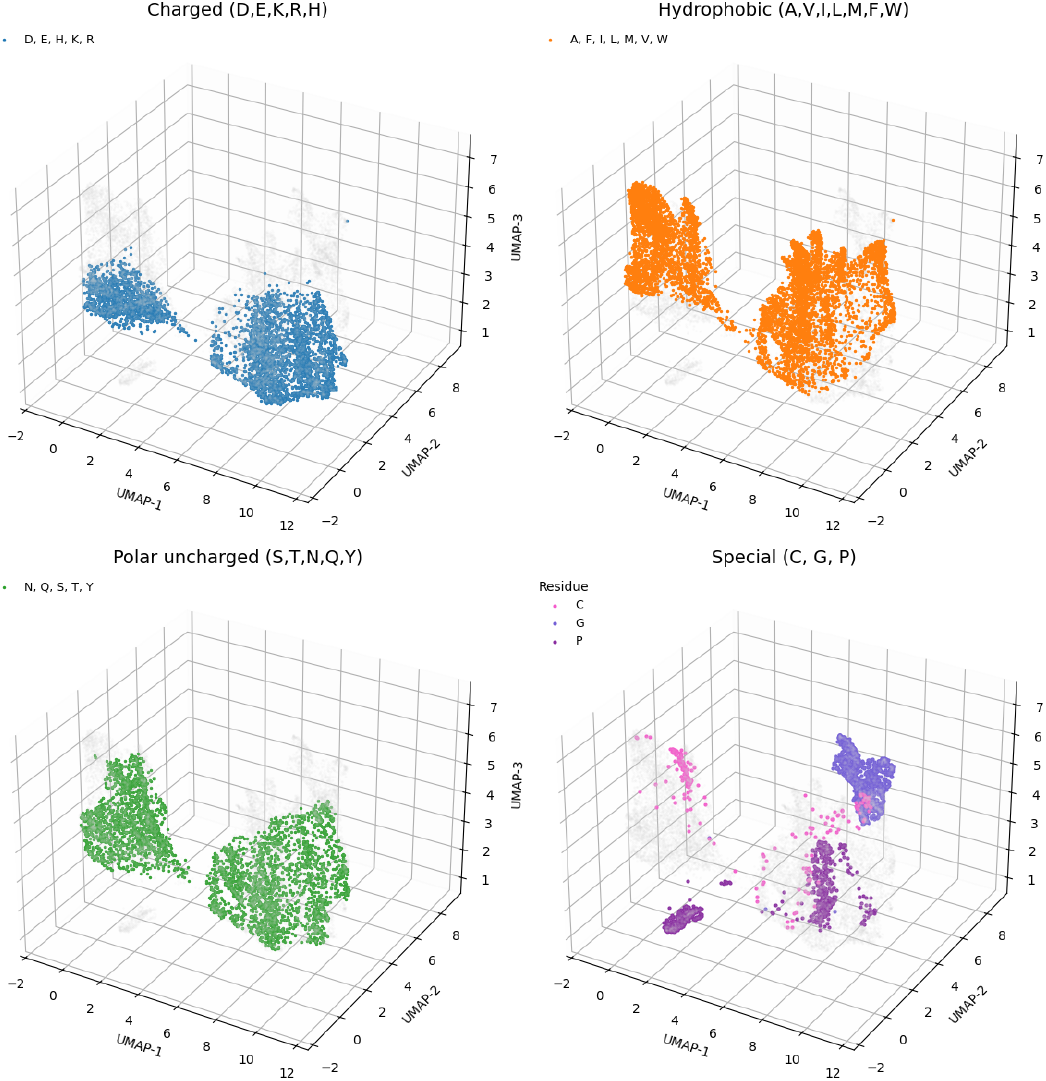
3D UMAP projection of CATH residue embeddings colored by amino acid type.

#### E.2.3 Structure ensemble analysis

##### Subsampled mdCATH

For each residue, we measure how much its embedding changes across the ensemble by averaging pairwise differences between frames. For a given residue and set of frames, we compute two physical descriptors: *Contact-map change*: we form a binary contact row per frame (contact if residues are within a chosen distance threshold) and measure, on average, what fraction of those contacts differ between frames. *Solvent-exposure change*: we compute solvent-accessible surface area (SASA), convert to residue-type–normalized relative SASA, and take the average absolute change between frames. We fit a simple linear model that predicts per-residue embedding change from the two descriptors. We aggregate performance on held-out residues and report: (i) the proportion of variance explained and (ii) the Spearman rank correlation between observed and predicted embedding change.

##### Rosetta Decoys

For each native protein we have a residue–embedding matrix and a set of its decoy matrices, aligned by residue index. We apply row-wise L2 normalization so that inner products equal cosine similarity. For a given protein, we compute the mean residue-wise cosine similarity between each decoy and its native, then take the average over decoys. The *native–decoy cosine margin* is defined as the difference between the native’s self-similarity (equal to 1.0 after normalization) and this mean decoy similarity.

To test linear separability at the residue level and generalization to unseen proteins, we train a logistic-regression classifier on residue embeddings with leave-protein-out grouped cross-validation: each residue embedding is a sample (label 0=native, 1=decoy) and carries its protein ID for grouped CV. We split with `GroupKFold` so all residues from a held-out protein appear only in the test set, and train an L2-regularized `LogisticRegression`. On each test fold we report AUROC; metrics are aggregated as mean *±* sd across folds.

### E.3 Per-residue generative model assessment

We compare distribution coverage of all-atom chemical environments sampled by generative models, stratified by residue type. For each residue type, we extracted the SLAE embeddings of 2000 random examples from the sequence-augmented CATH dataset and from a collection of 20,000 unconditional samples of all-atom protein structures from La-Proteina, Protpardelle-1c, and Chroma.

### E.4 Latent space interpolation

In Figure 9 A and B we show 20 out of 50 interpolated structures for AdK and KaiB. In addition, we compare linearly interpolated AdK structures from the SLAE latent space to those from the all-atom generative model Protpardelle-1c (Figure 10) and show that SLAE interpolation is better matched to simulated intermediate structures.

**Figure 8.**
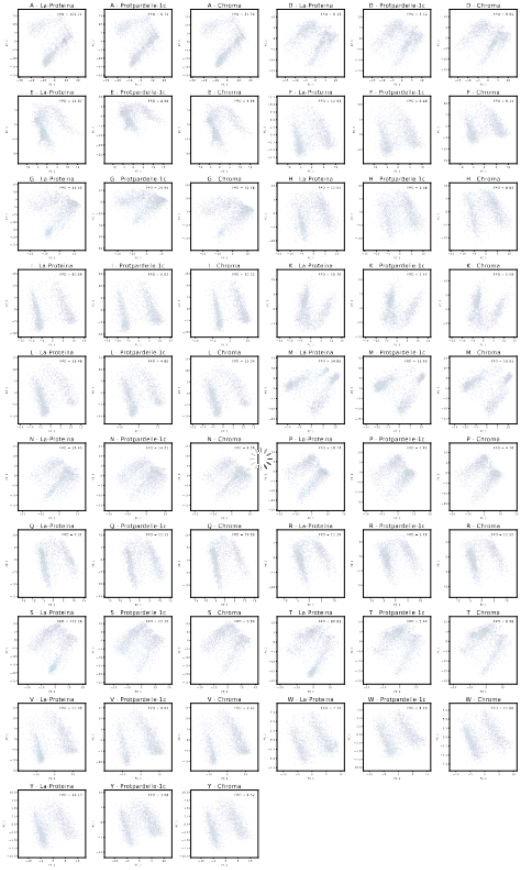
SLAE embeddings to assess residue environment coverage. PCA of SLAE per-residue embeddings of de novo structure samples (light blue) compared to the reference CATH distribution (purple) stratified by amino acid type given in the title. The two modes in each amino acid type correspond to residues belonging to a beta sheet or alpha helix.

**Figure 9.**
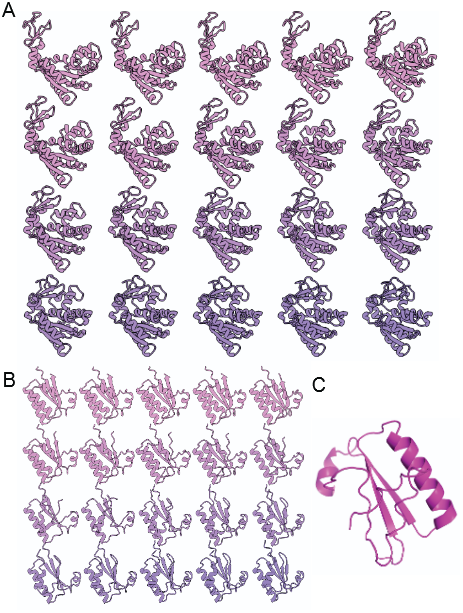
Structures decoded from SLAE latent interpolation. **A.** AdK **B**. KaiB **C**. Step 23 KaiB intermediate structure with under-characterized C-terminus showing disordered backbone collapsing onto itself.

**Figure 10.**
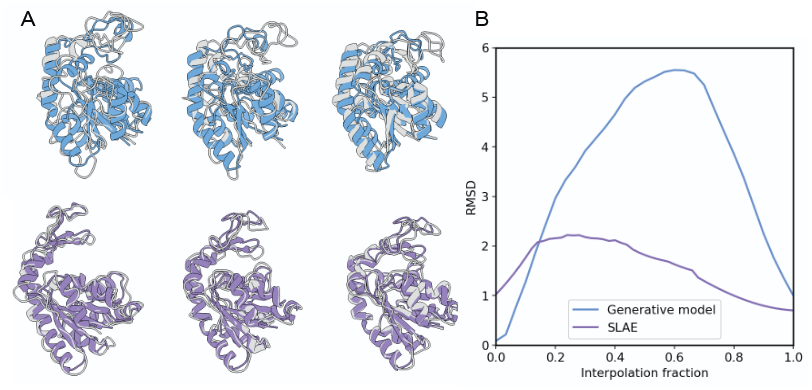
Comparison of SLAE and generative model (Protpardelle-1c) latent interpolation. **A.** Three representative steps from interpolation fraction 0.3 to 0.7. Top: Protpardelle-1c linear interpolation (blue) and best MD frame matches (grey). Bottom: SLAE linear interpolation (purple) and best MD frame matches (grey). **B**. RMSD of interpolation trajectories to their closest-match MD frames

## Footnotes

1 Fair comparison with open-sourced methods is not possible due to non-overlapping dataset splits (some entries from PLM-CS datasets do not pass the filter standard). We therefore re-trained a PLM-CS baseline on our splits and evaluate all embeddings under an identical protocol.

